# Dissociable Microstructural Correlates of Learning Rate and Learning Noise in Gamified Reward-Based Decision-Making

**DOI:** 10.64898/2026.02.24.707646

**Authors:** Melina Vejlø, Niia Nikolova, Leah Banellis, Ashley Tyrer, Vasilisa Skvortsova, Tobias U. Hauser, Micah G. Allen

**Affiliations:** Centre of Functionally Integrative Neuroscience, Aarhus University, Denmark; Department of Psychiatry and Psychotherapy, Faculty of Medicine, University of Tübingen, Tübingen, Germany; German Center for Mental Health (DZPG), Project Site Tübingen, Tübingen 72076, Germany; Max Planck UCL Centre for Computational Psychiatry and Ageing Research, University College London, London, UK; Functional Imaging Laboratory (FIL), Department of Imaging Neuroscience, University College London, London, UK; Cambridge Psychiatry, University of Cambridge, United Kingdom

**Keywords:** Reward Learning, Adaptive Behaviour, Voxel Based Quantification, MPM

## Abstract

Humans learn which actions yield the highest rewards through trial and error, gradually forming expectations about outcomes. Yet, people differ substantially in how quickly and precisely they learn. Such individual variability may partly be explained by differences in the brain’s microstructural organisation. In this large-scale study, 248 participants completed a gamified reward-learning task and underwent quantitative MRI to assess whole-brain microstructural indices of myelination (R1) and cortical iron (R2*). Using computational modelling, we quantified participants’ learning rates and learning noise, reflecting variability in how reward information is updated over time. Whole-brain voxel-based quantification analyses revealed that increased myelination in the cerebellum was associated with a higher learning rate, whereas learning noise was linked to increased myelination and iron concentration in the precentral gyrus. Together, these findings show that reward learning is not a unitary process but is instead shaped by distinct neurobiological pathways that support learning precision and noise. This work highlights how microstructural variation in sensorimotor and associative cortices contributes to stable versus variable reward learning behaviour across individuals.

**Significance Statement:** This study deepens our understanding of the neural microstructures involved in reward-related decision-making by employing advanced quantitative imaging and computational modelling to investigate brain microstructures associated with individual differences in noisy reward learning and decision-making, using a gamified task. Our results reveal distinct brain microstructures for learning rate and learning noise, which have not been reported before, thereby providing a new understanding of what may be involved in reward-based decision-making. These results provide new potential target pathways for future clinical research in psychiatric disorders where reward processing is implicated, such as ADHD and OCD.

## Introduction

Humans vary widely in their ability to integrate and learn from rewards, yet the biological origins of this variability remain poorly understood. Two key microstructural contributors may be particularly important. Firstly, iron is a critical factor in dopamine synthesis and regulation, and is thus likely to influence dopamine-related reward learning (Lozoff & Georgieff, 2006; Todorich et al., 2009). Moreover, cortical myelination is critical for information transmission and thus plays a key role in tasks that require information integration, such as reward learning (Grydeland et al., 2013; Guo et al., 2023; Samanez-Larkin et al., 2012; Xin & Chan, 2020). Variability in the availability of these microstructural contributors might explain why some individuals learn more quickly or reliably than others.

Canonical accounts of reinforcement learning (RL) suggest that surprising outcomes are learned by means of reward prediction errors, updating subjective representations of how good a choice option is. Traditionally, this learning process was thought to be unaffected by noise, and any variability in behaviour was attributed solely to a choice-selection stage, where stochasticity, biases, and exploratory tendencies (Dubois et al., 2021; Dubois & Hauser, 2022; Tervo et al., 2014; Wilson et al., 2014) come into play.

Recent work, however, suggests that learning itself also exhibits computational uncertainty and may be affected by noise (Drugowitsch et al., 2016; Findling & Wyart, 2021; Wyart & Koechlin, 2016). In their hallmark paper, Findling et al. (2019) demonstrated that more than half of seemingly stochastic choices could be explained by imprecision during learning. More recently, it has been suggested that the degree to which such learning noise affects behaviour differs substantially and meaningfully between individuals (Skvortsova & Hauser, 2022), and it has now been shown that adding computational noise in neural networks improves their performance (Findling & Wyart, 2024). Here, we speculate that this variation in learning distortions may arise in part from alterations in the brain’s microstructure.

Previous studies have mapped these different decision variables to neural activations in distinct brain regions supporting their computation. Trial-to-trial learning noise and prediction error have been linked to activity in the dorsal anterior cingulate cortex (dACC) (Behrens et al., 2007; Findling et al., 2019). Learning noise has also been associated with the dorsolateral prefrontal cortex (dlPFC), frontopolar cortex (FPC), and posterior parietal cortex (PPC) during outcome and choice periods, as well as negatively with ventromedial prefrontal cortex (vmPFC) activity during choice (Domenech et al., 2020; Findling et al., 2019). The understanding of the cerebellum’s role has evolved from being seen solely as involved in motor control to being recognised as a central hub that co-activates with frontal and parietal regions. (Stoodley, 2012). Recently, it has been linked specifically to reward learning and reward-related decision-making in both animals (Jin & Hull, 2025; Raymond, 2020) and humans (Nicholas et al., 2024). While functional MRI studies have mapped neural correlates of computational parameters, the structural correlates remain less explored. Voxel-Based Quantification (VBQ) is a promising new method for investigating individual differences, providing voxel-wise assessments of quantitative MRI maps (Draganski et al., 2011; Nikolova et al., 2025; Ziegler et al., 2019).

By using quantitative structural MRI to investigate both myelin- and iron-related microstructural integrity across 239 subjects, we show that distinct microstructural profiles underlie variability in learning efficiency and noise. Our analyses revealed that myelination in the cerebellum explained variability in learning rates, whereas myelination and iron concentrations in motor areas and the cerebellum drove differences in learning noise. By pairing quantitative MRI with computational modelling of behaviour, we were able to discover the microstructural constituents of learning variability.

## Materials & Methods

### Participants

565 participants (360 females, 205 males; median age = 24, age range = 18 - 56) participated in the Visceral Mind Project, a large-scale neuroimaging project at the Center of Functionally Integrative Neuroscience, Aarhus University. Participants were recruited through the SONA participation pool system and via local advertisements, including posters and social media. Inclusion criteria required participants to have corrected-to-normal vision and be fluent in either Danish or English. Participants did not take any medications except contraceptives or over-the-counter antihistamines. Furthermore, participants met standard MRI scanning requirements (i.e., no metal implants, claustrophobia, and not pregnant or breastfeeding). The participants from this dataset took part in multiple tasks, including MRI scans, physiological recordings both during scans and behaviour-only tasks, as well as psychiatric and lifestyle inventories, blood samples and computational tasks, spread over three visits on different days. The local Region Midtjylland Ethics Committee granted ethical approval for the study, and all participants provided written informed consent. The study was conducted in accordance with the Declaration of Helsinki (2013).

### Noisy reward learning task

To assess noisy reward learning, we used a gamified version of a two-armed bandit game, in which the bandits are depicted as brown and black-and-white space cows (Milky Way game on the Brain Explorer app, www.brainexplorer.net) (Skvortsova & Hauser, 2022). The goal of the game was to accumulate as many points as possible, which was translated into ‘space milk’. On each trial, participants made a self-paced choice between the two cows, after which the points earned were displayed. Participants were only shown the points from the chosen cow. Participants played 72 trials, split into two rounds of 36 trials each.

**Figure 1:**
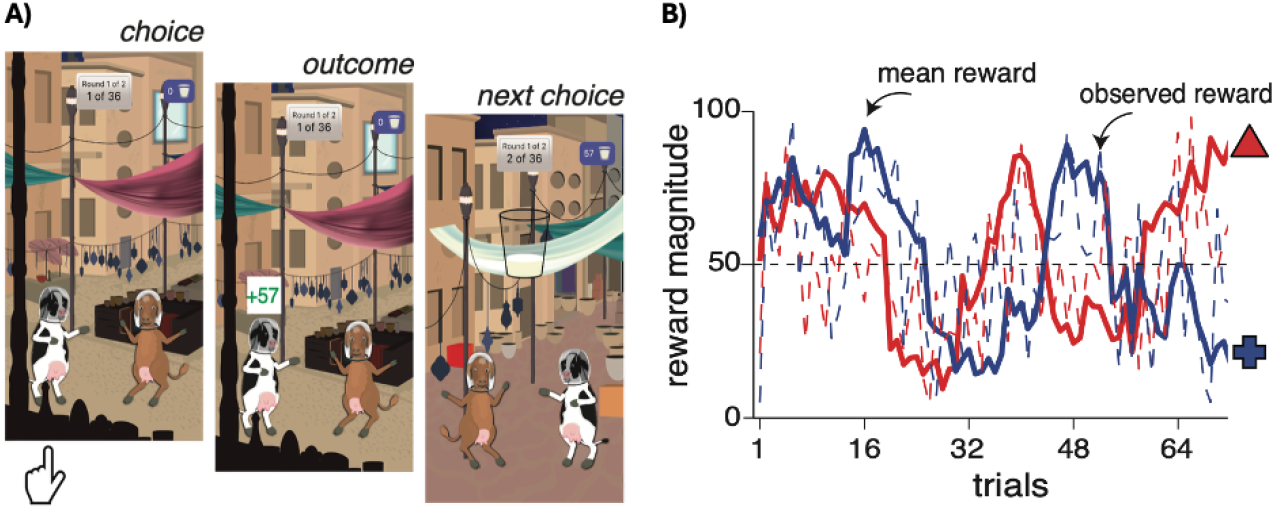
The trial structure of the Milky Way game. A) On each trial, participants were presented with two cows to choose between. The choice between the two is self-paced. After the choice, participants were presented with the outcome in the form of obtained points. This leads to the next choice. The points they won were represented in a cup of milk. Participants played two rounds, each consisting of 36 trials. Participants could keep track of the game’s progress at the top of the screen. B) Examples of reward walks (between 1 and 99 points) for the two bandits/cows. Reward magnitudes were sampled from two probability distributions with independently drifting means and changed throughout the trials. Both figures are from (Skvortsova & Hauser, 2022)

### Computational Model - Noisy learning model

Previous work has determined the best model for estimating this specific task (see Findling et al., 2019; Skvortsova & Hauser, 2022). To model participants’ choice behaviour, the same model that was previously developed and validated was used:

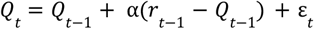

α is the learning rate, which scales the prediction error between the received reward, *r*_*t*−1_ and the expected reward, *Q*_*t*−1_. The updated values are influenced by ε_*t*_ which is an additive random noise drawn from a normal distribution with a zero mean and standard deviation σ_*t*_ equal to a constant fraction ζ of the magnitude of the prediction error: σ_*t*_ = ζ | *r*_*t*−1_ − *Q*_*t*−1_|.

In this noisy learning model, the noise added at each update scales with the prediction error in a way similar to Weber’s law. The learning noise was applied to the update of both chosen and unchosen option values. The choice process was modelled as a stochastic ‘softmax’ action selection policy, controlled by a ‘temperature’ β (choice stochasticity parameter) and an optional ‘choice repetition bias ξ:

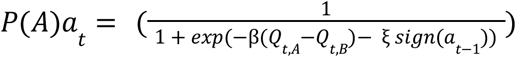

*Q*_*t,A*_ and *Q*_*t,B*_ are values associated with options A and B at time point *t*.

### Multi-Parameter Brain Mapping

We used a well-established qMRI protocol (Weiskopf et al., 2013, 2015) to map percentage saturation due to magnetisation transfer (MT), longitudinal relaxation rate (R1) and effective transverse relaxation rate (R2*), followed by voxel-based quantification to relate individual differences in reward learning to brain microstructure. The process of data acquisition and map creation has previously been described in Nikolova et al. (2025).

#### Data Acquisition

The imaging data were obtained with a 3T MR system (Magnetom Prisma, Siemens Healthcare, Erlangen, Germany), employing a standard 32-channel RF head coil and a body coil. High-resolution, whole-brain T1-weighted anatomical images (0.8 mm^3^ isotropic resolution) were captured using an MP-RAGE sequence, with a repetition time of 2.2 s, echo time of 2.51 ms, a matrix size of 256 × 256 × 192 voxels, a flip angle of 8°, and acquired in the anterior-posterior direction. Whole-brain images at 0.8 mm isotropic resolution were captured using a multi-parameter mapping (MPM) quantitative imaging protocol (Callaghan et al., 2019; Weiskopf et al., 2013). The sequence included three spoiled multi-echo 3D fast low-angle shot (FLASH) scans and three calibration scans to correct RF transmit field inhomogeneities. These FLASH scans were performed with MT, PD, and T1 weighting. The flip angles were set to 6° for MT- and PD-weighted images and 21° for T1-weighted images. MT-weighting employed a Gaussian RF pulse, 2 kHz off resonance, lasting 4 ms, with a nominal flip rate of 220°. The field of view measured 256 mm (head-foot), 224 mm (anterior-posterior), and 179 mm (right-left). Gradient echoes were collected with alternating readout gradient polarity at echo times from 2.34 to 13.8 ms (MT) or 18.4 ms (PD and T1), using a readout bandwidth of 490 Hz/pixel. For the MT scan, six echoes were acquired to maintain a TR of 25 ms across all FLASH volumes. Accelerated data collection was achieved with parallel imaging using the GRAPPA algorithm, with an acceleration factor of 2 in each phase-encoding direction and 40 reference lines. All acquisitions included a 30° slab rotation. B1 mapping involved 11 measurements with flips ranging from 115° to 65° in 5° steps. The entire quantitative MRI scanning process took approximately 26 minutes.

### Map Creation

All qMRI images were preprocessed with the hMRI toolbox v. 0.5.0 (January 2023) (Tabelow et al., 2019) and SMP12 (version 12.r7771, Wellcome Trust Centre for Neuroimaging, http://www.fil.ion.ucl.ac.uk/spm/) to correct for spatial transmit and receive field inhomogeneities and to generate quantitative maps of MT, PD, R1, and R2*. The toolbox was set up with standard settings, aside from enabling correction for imperfect spoiling. All images were first aligned to MNI standard space before creating the maps. Four maps were produced to model different tissue microstructure aspects: an MT map indicating myeloarchitectural integrity (Helms et al., 2008), a PD map showing tissue water content, an R1 map reflecting myelination, iron levels, and water content (mainly influenced by myelination) (Lutti et al., 2014), and an R2* map sensitive to tissue iron (Langkammer et al., 2010). The segmentation approach (Ashburner & Friston, 2005) was used to divide MT saturation maps into grey matter (GM), white matter (WM), and cerebrospinal fluid (CSF) probability maps. Tissue probability maps, derived from multi-parametric maps by Lorio et al. (Lorio et al., 2016), were used without bias field correction, as MT maps do not show significant bias field effects. These GM and WM maps facilitated inter-subject registration via Diffeomorphic Image Registration (DARTEL), a nonlinear diffeomorphic algorithm (Ashburner, 2007). The MT, PD, R1, and R2* maps were normalised to MNI space (1 mm isotropic resolution) using the DARTEL template and participant-specific deformation fields. Registration relied on MT maps due to their high contrast in subcortical regions and similar WM-GM contrast to T1-weighted images (Helms et al., 2009). Tissue-weighted smoothing with a 4 mm FWHM kernel was applied via the VBQ method (Draganski et al., 2011). Unlike voxel-based morphometry, VBQ minimises partial volume effects and preserves original quantitative data by not modulating parameter maps for volume. The GM segments from each map served for all statistical analyses.

### Participant Exclusion Criteria and MRI Quality Control

Following inspection, several participants were excluded due to MRI or behavioural issues. Three participants were removed immediately after MR data collection for medical reasons (one cerebral palsy, two suspected brain abnormalities). MPM contrast images were obtained for 503 participants, with three excluded due to scanning errors. Because MPM data is sensitive to motion artefacts, a thorough quality control (QC) was performed using visual HTML reports from the hMRI-vQC toolbox, reviewed by two researchers. Based on QC, 57 participants were excluded due to excessive motion affecting tissue-class segmentation. Data from 443 participants were used in the spatial analysis and template creation with DARTEL. Additional quality assessments with CAT12 (Gaser et al., 2024) led to further exclusions: 19 from MT maps, 20 from R1 maps, and 24 from R2* maps. A total of 311 participants completed the reward learning task. In total, we had MT maps and task data from 239 participants for VBQ analyses (165 females, 74 males, median age =24.3+-4), further subsuming into 236 for R1 (166 females, 70 males, median age = 24.4+-4.2), and 232 for R2* (165 females, 67 males, median age = 24.2+-3.7).

## Data Availability

Code, demographics data, and anonymised summary variables are publicly accessible on GitHub (https://github.com/embodied-computation-group/noisy_reward_learning_VBQ). Summary statistical brain maps can be found on NeuroVault (https://neurovault.org/collections/HSOJRRQV/). The neuroimaging data presented in this study are part of a larger dataset and will be released publicly in a dedicated paper after the mandatory data embargo, which ensures participants’ data privacy rights.

### Voxel-Based Quantification Analysis

Grey and white matter masks were generated based on our samples by averaging the smoothed, modulated GM and WM segment images, and thresholding the result at p >.2. Inter-subject variation in MT, R1 and R2* GM maps was modelled in separate multiple linear regression analyses. The learning rate for the chosen and unchosen object, zeta (learning noise) and tau (Choice Stochasticity) were used as regressors of interest in the VBQ analysis. We also included age, gender, body mass index (BMI) and total intracranial volume (TIV) as nuisance covariates in all analyses, following recommended procedures for computational neuroanatomy (Ridgway et al., 2008).

All statistical analyses were conducted in SPM12. Threshold-Free Cluster Enhancement (TFCE) was applied to the resulting statistical maps using the TFCE toolbox (Smith & Nichols, 2009) for SPM12 (Wellcome Centre for Human Neuroimaging, London, UK). TFCE was performed with 5,000 permutations to derive corrected p-values under the null distribution. Default TFCE parameters were used (E = 0.5, H = 2.0). Statistical significance was defined as p <.05, with a family-wise error (FWE) cluster correction. All analyses were confined to a whole-brain grey matter mask.

## Results

To examine the relationship between computational parameters (learning rates, choice stochasticity, and learning noise) derived from participant behaviour and indices of cortical microstructure, we employed whole-brain, TFCE-corrected multiple linear regression. This enabled us to estimate the microstructural correlates of noisy reward learning.

### Increased myelination in the right cerebellum is associated with learning rate and learning noise

We found that R1 values, associated with myelin density, in the left cerebellum exterior (k = 35385, pFWEcorr =.002, peak voxel coordinates: x = -16.8, y = -52, z = -23.2) were positively associated with the learning rate for the chosen object. We found no significant cluster for the learning rate for the unchosen object. This finding suggests a positive association between learning about rewards and increased myelination density in the right cerebellum, consistent with previous findings that reward learning is associated with cerebellar activation (Nicholas et al., 2024).

**Figure 2:**
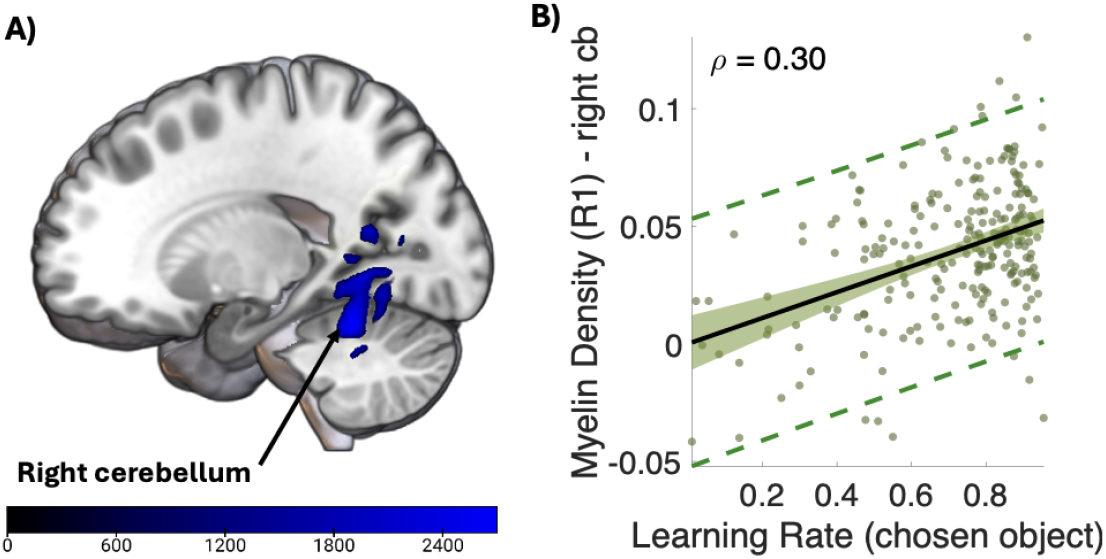
Microstructural correlates for learning rate (α) for the chosen object in R1. A) R1 values in the right cerebellum exterior were positively associated with the learning rate for the chosen object. Colourbar shows the TFCE value. B) The learning rate for the chosen object (on the x-axis) was positively correlated with R1 values in the right cerebellum exterior (y-axis). Maps were TFCE (non-parametric) cluster corrected and FWE corrected for multiple comparisons (pFWE <.05). Scatter plot is for visualisation purposes only.

Moreover, we found that R1 values in the right cerebellum (k = 24875, pFWEcorr =.012, peak voxel coordination: x = 14.5, y = -58.4, z = -28.8) were positively associated with learning noise. We found no significant clusters for choice stochasticity for myelin density. This indicates that learning noise, but not choice stochasticity, is related to an increase in myelin density in the right cerebellum.

**Figure 3:**
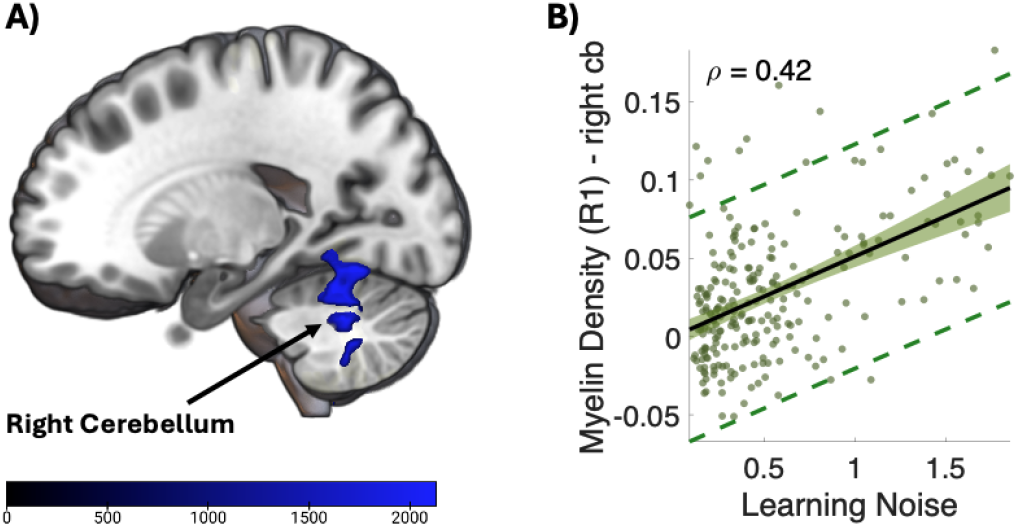
Microstructural correlates for learning noise and R1. A) R1 values in the right cerebellum were positively associated with learning noise values. The colourbar indicates the TFCE corrected value. B) R1 values on the y-axis in the right cerebellum were positively associated with the learning noise on the x-axis. Maps were TFCE (non-parametric) cluster corrected, and FWE corrected for multiple comparisons (pFWE <.05). Scatter plot is for visualisation purposes only.

### Myelin density and iron concentration in the left medial precentral gyrus are associated with learning noise

We found that R2* values, associated with iron concentration, in the left precentral gyrus medial segment (k = 105, pFWEcorr =.04, peak voxel coordinates: x = -3.2, y = -22.4, z = 71.2) were positively associated with learning noise. The same is the case for R1 values in the left medial segment of the precentral gyrus (k = 1073, pFWEcorr =.016, peak voxel coordinates: x = -3.2, y = -24, z = 72.8). The two clusters overlap, indicating increased myelination and iron concentration in the same brain area. We found no association between choice stochasticity and iron concentration. These findings indicate that both increased myelin density and iron concentration are associated with learning noise in the motor cortex.

**Figure 4:**
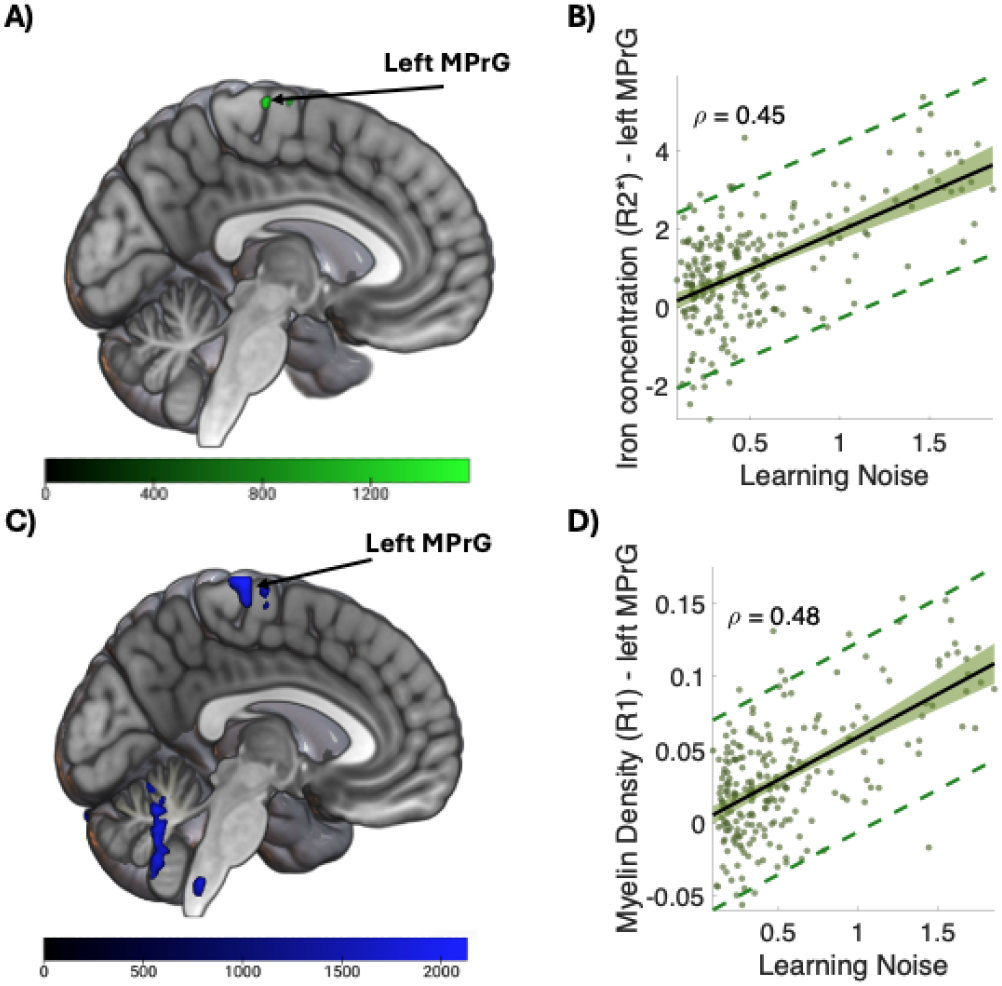
Microstructural correlates of learning noise for iron concentration (R2*) and myelin density (R1) A) R2* values in the left precentral gyrus medial segment were positively associated with the learning noise. B) R2* values (y-axis) in the left precentral gyrus medial segment were positively associated with the learning noise (x-axis). C) R1 values in the medial segment of the precentral gyrus were positively associated with learning noise values. The colourbar indicates the TFCE corrected value. D) R1 values on the y-axis in the left precentral gyrus medial segment were positively associated with the learning noise on the x-axis. Maps were TFCE (non-parametric) cluster corrected, and FWE corrected for multiple comparisons (pFWE <.05). Scatter plot is for visualisation purposes only.

For the complete list of significant clusters, see Table 2-4 in the supplementary information. We found no significant associations between model parameters and MT map values, indicating no obvious relationship between computational parameters of noisy reward learning and myeloarchitectural integrity.

## Discussion

In this study, we investigated the microstructural brain correlates of noisy reward learning by combining VBQ with computational modelling of behaviour in a healthy adult cohort. Using whole-brain, TFCE-corrected analyses, we found that the learning rate for the chosen object and learning noise was associated with increased R1 values, indicating an increase in myelin density, in the cerebellum, while learning noise was associated with R2*, indicative of iron concentration, and R1 values in the medial precentral gyrus. These findings offer new insights that variability in reinforcement learning parameters is associated with regional microstructural differences in both cortical and subcortical regions, particularly concerning myelin density and iron concentration.

VBQ offers a promising method for investigating individual differences by providing voxel-wise assessment of quantitative MRI maps (Draganski et al., 2011). Unlike traditional voxel-based morphometry, VBQ relies on biophysically meaningful parameters, reducing interpretational ambiguity and enabling inferences about more specific tissue properties. Moreover, quantitative contrasts such as R2* are sensitive to tissue iron, which accumulates in neuromelanin-containing dopaminergic neurons and thereby serves as an indirect marker of dopamine system integrity (Stüber et al., 2014). This further links quantitative MRI markers to neurochemical systems central to reinforcement learning (Depierreux et al., 2021).

The association between increased myelination in the motor cortex and learning noise has not been previously reported. The motor cortex has been primarily implicated in motor skill learning, with fMRI studies showing its involvement in motor learning tasks and in reward paradigms that include a motor component (e.g., response selection) (Hamano et al., 2021). Even in ostensibly cognitive tasks, updating action values often requires motor responses, such as choosing between left and right. If learning noise reflects imprecision in value updating (Findling et al., 2019), these imprecisions may propagate into the motor planning and selection system. The motor system may therefore actively integrate expected values of actions, and higher myelination may influence the efficiency or adaptability of motor computations, accounting for the observed link with learning noise. Animal studies support this interpretation: in rodents trained on skilled, reward-based motor tasks, increased myelination in white matter under the motor cortex has been observed histologically and shown to correlate with learning rate (Sampaio-Baptista et al., 2013). Moreover, detailed imaging studies in mice demonstrate that motor learning drives dynamic myelin remodelling, targeted to learning-activated axons, within the primary motor cortex, in association with behavioural improvements (Bacmeister et al., 2022).

We also found that the learning rate for the chosen object was associated with increased cerebellar myelination. While traditionally linked to motor control, the cerebellum is now recognised as a hub engaged across multiple cognitive domains, including working memory, language, and spatial processing (Jacobi et al., 2021). Cerebellar activation during cognitive tasks often co-occurs with activity in prefrontal and parietal regions (Stoodley, 2012), and its role in forming internal models of action (Ito, 2008) may align with the observed relationship between cerebellar myelination and reinforcement learning rates. More recently, the cerebellum has been linked more directly to reward learning, specifically in patients with ataxia, who show decreased trial-to-trial reward learning compared with healthy controls (Nicholas et al., 2024).

These findings extend prior work on functional correlates of reinforcement learning parameters (Hauser et al., 2017; Skvortsova & Hauser, 2022) by demonstrating associations with structural markers of myelination and iron. One intriguing source of variability is nutritional status. Nutritional iron deficiency reduces myelination and can influence neuromelanin-containing structures, affecting R2* and related MRI contrasts (Todorich et al., 2009), potentially contributing to individual differences in reinforcement learning. Conversely, iron accumulation is known to occur with ageing and can degrade myelin integrity (Khattar et al., 2021; Steiger et al., 2016). Steiger et al. (2016) showed that an increased iron-to-myelin ratio in the ventral striatum predicted memory performance in older adults, underscoring the importance of iron–myelin interactions for learning-related processes. Our results suggest that similar mechanisms may extend to reinforcement learning and decision-making.

The role of dopamine also warrants further investigation. Iron content and neuromelanin-sensitive MRI contrasts such as R2* are linked to the integrity of dopaminergic neurons and can serve as indirect markers of dopaminergic system structure and function (Depierreux et al., 2021), suggesting that structural measures may provide indirect proxies of dopaminergic tone. This raises the possibility that inter-individual differences in learning noise and learning rate are partly explained by dopaminergic variability. Future work combining quantitative MRI with positron emission tomography (PET) could test this more directly by linking dopamine receptor density with microstructural correlates of reinforcement learning.

Despite these insights, the present study has limitations. The correlational design does not establish causal pathways between brain structure, computational parameters, and behaviour. Longitudinal studies are needed to disentangle whether variability in myelination, iron, and neuromelanin reflects stable individual differences, developmental factors, or modifiable states such as nutrition. The surprising association of motor cortex myelination and learning noise, in particular, requires replication and deeper mechanistic investigation. Furthermore, our findings motivate future work in clinical populations, such as ADHD and OCD, where non-greedy decisions have been differentially linked to learning noise and choice stochasticity (Skvortsova & Hauser, 2022). Examining whether microstructural variation in these groups underlies distinct patterns of non-greedy choice could clarify the biological bases of impulsivity and compulsivity.

In sum, this study provides novel evidence linking reinforcement learning parameters to brain microstructure, highlighting the cerebellum and motor cortex. These results open new directions for understanding how variability in myelination, iron, and neuromelanin contributes to adaptive decision-making, and point toward integrative imaging approaches that combine quantitative MRI, PET, and longitudinal designs to clarify the structural underpinnings of reward learning.

## Acknowledgements

This research is financially supported by a Lundbeckfonden Fellowship (R272-2017-4345) and a European Research Council Grant (ERC-2020-StG-948788) awarded to MGA. MV, LB, and AT were supported by ERC-2020-StG-948788. The funding sources were not involved in the study design, collection, analysis, interpretation, or writing of the manuscript. TUH has received funding from the Wellcome Trust (316955/Z/24/Z), the European Research Council (ERC) under the European Union’s Horizon 2020 research and innovation programme (grant agreement No 946055), and the Carl-Zeiss-Stiftung. This work was supported by the Alexander von Humboldt foundation, (more) precisely the Alexander-von-Humboldt-Professorship award to Peter Dayan. For the purpose of Open Access, the author has applied a CC BY public copyright license to any Author Accepted Manuscript version arising from this submission.

While preparing the manuscript, the authors used AI-assisted language editing tools to improve grammar, clarity, and readability. These tools were used only for language refinement and did not contribute to the study design, data analysis or scientific conclusions. The authors carefully reviewed and approved all content and take full responsibility for the manuscript.

