## Supplemental Information for "Dissociable Microstructural Correlates of Learning Rate and Learning Noise in Gamified Reward-Based Decision-Making"

**Supplementary figures and tables**

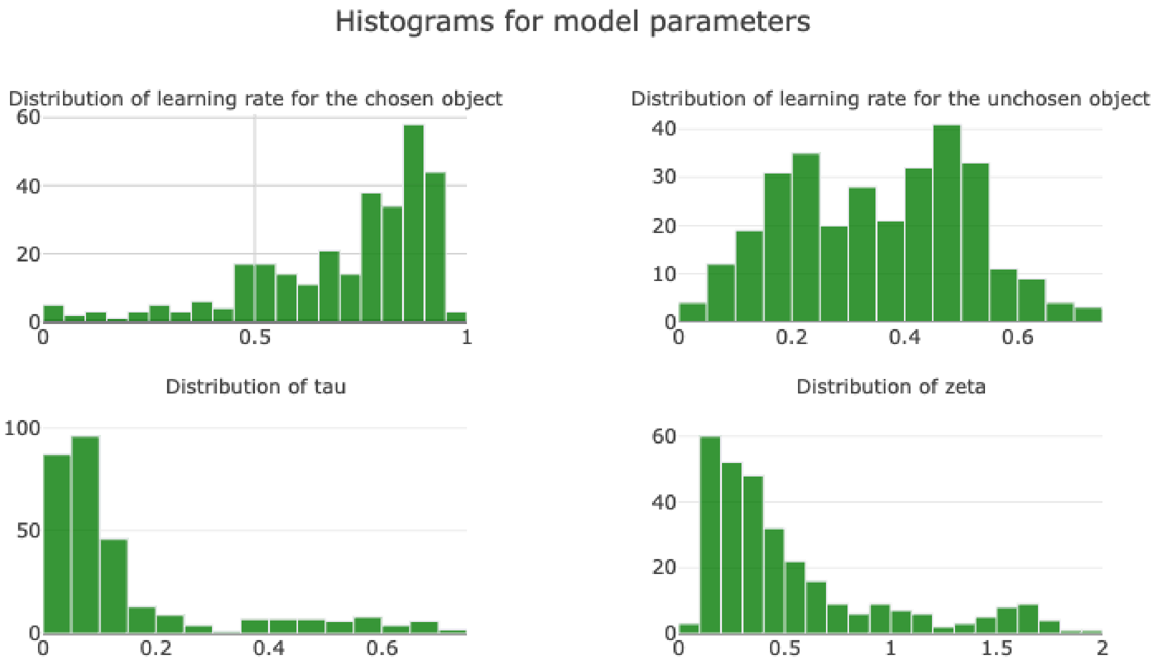


***Figure S1: Histogram of model parameters***

Histograms for the four computational parameters used for VBQ analysis, learning rates for the chosen and unchosen objects (cows), choice stochasticity (tau), and learning noise (zeta).

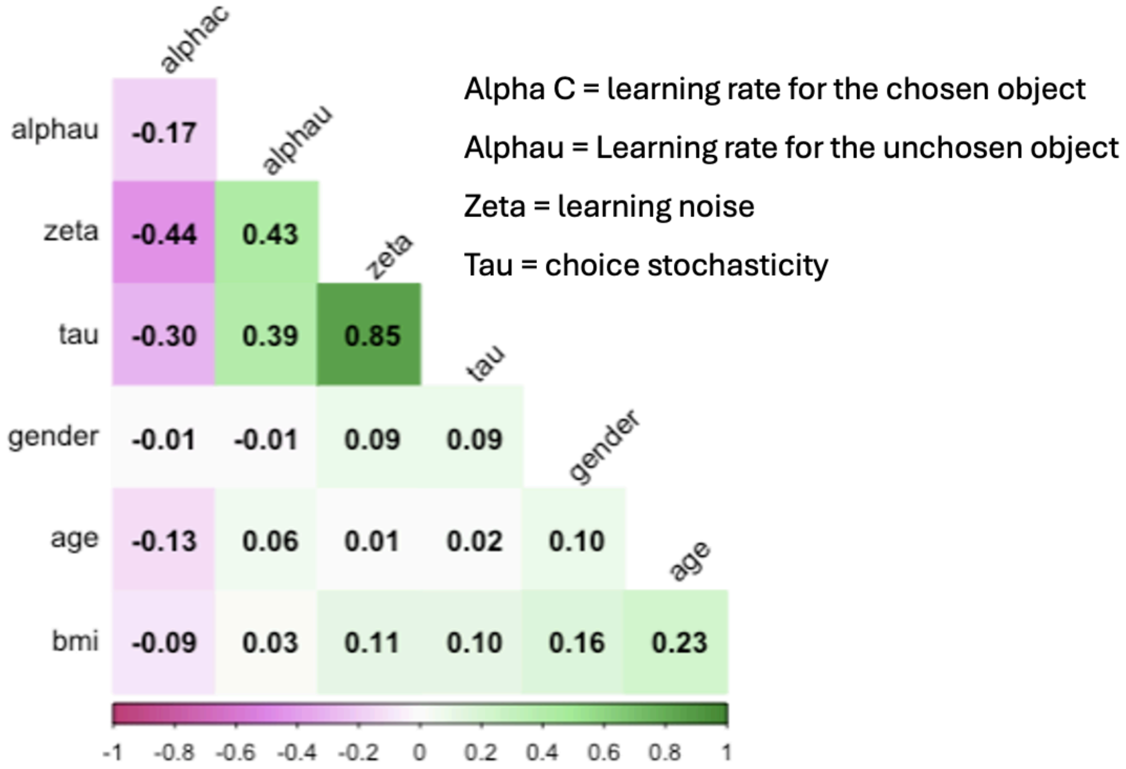


***Figure S2: Pairwise correlations between model parameters and confounding factors.***

**Table 1: Participant Demographics of 311 Noisy reward learning participants**

| Variable | Mean (± SD) |
|---|---|
| Age | 24.6 ± 4.4 |
| Gender | 208 (68.6%) female, 95 (31.4%) male |
| BMI | 22.5 ± 2.8 |

**Table 2: Summary of Whole Brain VBQ Results: Positive correlation between R1 and zeta (learning noise)**

| | Significant Whole Brain Clusters | | | | | |
|---|---|---|---|---|---|---|
| **Region** | ***k*** | ***p*(FWE-corr)** | **TFCE** | **x** | **y** | **z** |
| R Cerebellum Exterior | 24875 | 0.012 | 2131.6 | 14.4 | -58.4 | -28.8 |
| L Precentral Gyrus Medial Segment | 1073 | 0.016 | 2012.76 | -3.2 | -24 | 72.8 |
| R supplementary Motor Cotex | 162 | 0.04 | 1656.75 | 3.2 | -13.6 | 58.4 |
| R Precentral Gyrus | 201 | 0.045 | 1605.61 | 22.4 | -24 | 65.6 |
| L Cerebral White Matter | 202 | 0.046 | 1597.49 | -5.6 | -14.4 | 69.6 |
| R Cerebellum Exterior | 183 | 0.049 | 1591.39 | 12 | -74.4 | 53.6 |
| L Supplementary Motor Cortex | 9 | 0.049 | 1582.35 | 0 | -13.6 | 59.2 |
| L Cerebellum Exterior | 20 | 0.049 | 1571.99 | -36 | -53-.6 | -41.6 |

**Table 3: Summary of Whole Brain VBQ Results: Positive correlation between R1 and alpha C (learning rate for the chosen object)**

| Significant Whole Brain Clusters | | | | | | |
|---|---|---|---|---|---|---|
| **Region** | ***k*** | ***p*(FWE-corr)** | **TFCE** | **x** | **y** | **z** |
| L Cerebellum Exterior | 35385 | 0.002 | 2700.03 | -16.8 | -52 | -23.2 |
| L Cerebellum White Matter | 1 | 0.034 | 1644.71 | -9.6 | -46.4 | -28.8 |
| L Calccarine Cortex | 590 | 0.035 | 1631.98 | -4 | -72 | 12.8 |
| R Calcarine Cortex | 759 | 0.038 | 1605.39 | 19.2 | -66.4 | 7.2 |
| L Interior Temporal Gyrus | 264 | 0.04 | 1585.91 | -49.6 | -65.6 | -13.6 |
| R precuneus | 2 | 0.042 | 1564.75 | 20 | -51.2 | 8.8 |
| L Calcarine Cortex | 27 | 0.05 | 1508.94 | -17.6 | -70.4 | 8.8 |

**Table 4: Summary of Whole Brain VBQ Results: Positive correlation between R2* and zeta (learning noise)**

| Significant Whole Brain Clusters | | | | | | |
|---|---|---|---|---|---|---|
| **Region** | ***k*** | ***p*(FWE-corr)** | **TFCE** | **x** | **y** | **z** |
| L Precentral Gyrus medial segment | 105 | 0.04 | 1564.77 | -3.2 | -22.4 | 71.2 |
| L Cerebral White Matter | 36 | 0.048 | 1503.95 | -6.4 | -12.8 | 70.4 |
